# SARS-CoV-2 introduction and lineage dynamics across three epidemic peaks in Southern Brazil: massive spread of P.1

**DOI:** 10.1101/2021.07.29.454323

**Authors:** Ana Paula Muterle Varela, Janira Prichula, Fabiana Quoos Mayer, Richard Steiner Salvato, Fernando Hayashi Sant’Anna, Tatiana Schäffer Gregianini, Letícia Garay Martins, Adriana Seixas, Ana B. G. Veiga

**Author notes:** **Corresponding author:** A. B. G. Veiga. Departamento de Ciências Básicas da Saúde, Universidade Federal de Ciências da Saúde de Porto Alegre. Rua Sarmento Leite, 245. CEP 90050-170. Porto Alegre, RS, Brazil. **Note:** APMV and JP contributed equally in generating data.

## Abstract

**Background:** Genomic surveillance of SARS-CoV-2 is paramount for understanding viral dynamics, contributing to disease control. This study analyzed SARS-CoV-2 genomic diversity in Rio Grande do Sul (RS), Brazil, including the first case of each Regional Health Coordination and cases from three epidemic peaks.

**Methods:** Ninety SARS-CoV-2 genomes from RS were sequenced and analyzed against SARS-CoV-2 datasets available in GISAID for phylogenetic inference and mutation analysis.

**Results:** SARS-CoV-2 lineages among the first cases in RS were B.1 (33.3%), B.1.1.28 (26.7%), B.1.1 (13.3%), B.1.1.33 (10.0%), and A (6.7%), evidencing SARS-CoV-2 introduction by both international origin and community-driven transmission. We found predominance of B.1.1.33 (50.0%) and B.1.1.28 (35.0%) during the first epidemic peak (July–August, 2020), emergence of P.2 (55.6%) in the second peak (November–December, 2020), and massive spread of P.1 and related sequences (78.4%), such as P.1-like-II, P.1.1 and P.1.2 in the third peak (February–April, 2021). Eighteen novel mutation combinations were found among P.1 genomes, and 22 different spike mutations and/or deletions among P.1 and related sequences.

**Conclusions:** This study shows the dispersion of SARS-CoV-2 lineages in Southern Brazil, and describes SARS-CoV-2 diversity during three epidemic peaks, highlighting the spread of P.1 and the high genetic diversity of currently circulating lineages. Genomic monitoring of SARS-CoV-2 is essential to guide health authorities’ decisions to control COVID-19 in Brazil.

**Summary:** Ninety SARS-CoV-2 genomes from Rio Grande do Sul, Brazil, were sequenced, including the first cases from 15 State Health Coordination regions and samples from three epidemic peaks. Phylogenomic inferences showed SARS-CoV-2 lineages spread, revealing its genomic diversity.

## Introduction

The new coronavirus SARS-CoV-2 emerged at the end of 2019 in the province of Wuhan, China, and rapidly spread to other countries, infecting millions of people worldwide [1,2]. In Brazil, as of 27 July 2021 approximately 19.7 million cumulative cases and 550,000 deaths have been reported, ranking third country in the cumulative number of cases and second in the number of deaths [3]. The first confirmed case in Brazil was a man returning from Italy to São Paulo on 26 February 2020; following that, different viral strains were introduced in the country by individuals returning from international travel [4]. Notably, with high viral transmission, variants of concern (VOC), variants of interest (VOI) and variants under investigation (VUI) emerged in the Brazilian population, such as P.1, P.2 and VUI-NP13L, respectively, all derived from B.1.1.28 [5–8].

Rio Grande do Sul (RS), the southernmost Brazilian state, reports high numbers of respiratory viral infections annually [9]. RS is a hub for connecting flights, with four international airports and 56 regional airports, seven of which have regular routes; moreover, RS borders Uruguay and Argentina, is close to Paraguay, and is a popular tourist destination in Brazil [10]. Hence, RS is one of the Brazilian states with the highest number of reported SARS-CoV-2 variants [11].

In RS, there are 19 Regional Health Coordinations (RHC) responsible for epidemiological surveillance, whose data show three COVID-19 epidemic peaks in the state until June 2021 [11]. The first peak occurred in August 2020, when the cumulative number of cases per day was around 3,000; the second peak was in November–December 2020, with approximately 6,000 cases reported daily; the third and more critical peak started in February and lasted until April 2021, in which more than 13,000 cases were reported in a single day (**Figure 1**). Notably, in March 2021 RS became the epicenter of COVID-19 in Brazil, with great health and economic burden, and exhaustion of the health system [11].

**Figure 1.**
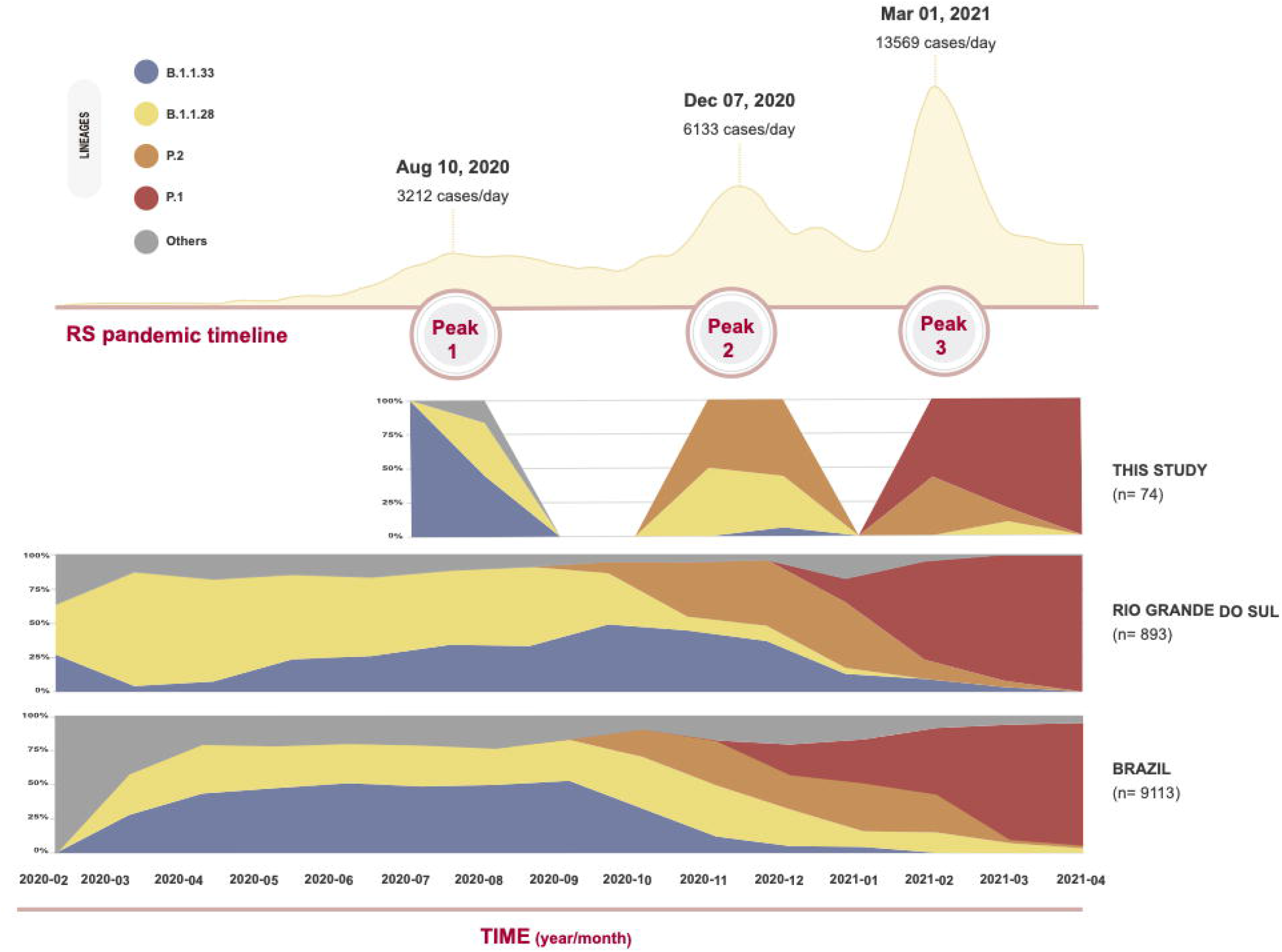
Profile of SARS-CoV-2 lineage distribution from March 2020 to April 2021. Cumulative number of COVID-19 confirmed cases in RS state during three peaks (July–August, 2020; November–December, 2020; and February–April, 2021), and density distribution of the main lineages identified in this study.

To better understand the spread of SARS-CoV-2 lineages in RS, we performed genomic analysis of SARS-CoV-2 from the first cases of 15 RHCs, in addition to samples from three epidemic peaks. Our findings help to understand SARS-CoV-2 transmission and diversity in Southern Brazil, contributing to the Brazilian Public Healthcare System.

## Methods

### Study design

Ninety upper- and lower-respiratory tract secretion samples from patients with SARS-CoV-2 infection confirmed by RT-qPCR were obtained from the State Central Laboratory of Rio Grande do Sul (LACEN-RS) (**Supplementary Data 1**). To investigate spread and diversity of SARS-CoV-2 lineages across RS we selected the first positive case from 15 RHCs of RS (n = 15) (**Supplementary Table 1**), and samples (n = 75) from the four cities with the highest cumulative incidences (Santo Angelo, Passo Fundo, Caxias do Sul, and Porto Alegre), comprising three epidemic peaks: July–August 2020 (first peak; n = 20), November–December 2020 (second peak; n = 18), and February–April 2021 (third peak; n = 37) (**Figure 1**). This study was approved by the Ethics Committee of Universidade Federal de Ciências da Saúde de Porto Alegre (Protocol n. 3.978.647, CAAE 30714520.0.0000.5345).

### RNA extraction and sequencing

Viral RNA was extracted with MagMAX™ Viral/Pathogen Nucleic Acid Isolation kit in a KingFisher™ Flex Purification System (ThermoFisher Scientific). SARS-CoV-2 RT-qPCR was performed in a 7500 Real-Time PCR System (Applied Biosystems) using Seegene-Allplex 2019-nCoV Assay, following the manufacturer’s instructions.

Libraries were constructed using QIASEQ SARS-CoV-2 Primer Panel and QIAseq FX DNA Library UDI-A kit (Qiagen) following manufacturer’s instructions, with previously described annealing temperature [12]. Libraries were quantified using QubitTM dsDNA HS Assay kit and normalized to equimolar concentrations. Sequencing was performed with MiSeq Reagent Kit v3 600 cycles in a MiSeq instrument (Illumina, USA).

### Bioinformatic analysis

Raw fastq files were trimmed to remove adapter sequences and low-quality reads using trimmomatic [13] (Parameters: leading:3 trailing:3 slidingwindow:4:25 minlen:36 illuminaclip: TruSeq3-PE.fa:2:30:10). Reads were mapped against SARS-CoV-2 reference genome (GenBank Accession NC_045512.2) using the bwa-mem algorithm (v0.7.17) [14] under default parameters. The quality of the resulting mapped BAM files was checked using Qualimap [15]. The software iVar (v1.2.2) was used to trim QIASEQ SARS-CoV-2 Primer Panel sequences and along with Samtools (v1.6) mpileup function [16] to variant calling (parameters: -A -d 20000 -Q 0) and consensus sequence generation (parameters: -q 20 -t 0). Consensus sequences were submitted to Nextclade (https://clades.nextstrain.org) for quality inspection and to determine the mutational profile. Sequences were submitted to Phylogenetic Assignment of Named Global Outbreak Lineages (Pangolin) v2.4.2 (https://github.com/cov-lineages/pangolin) for lineage assignment. Mutations related to lineage P.1 were also manually inspected for assessment of lineages P.1.1, P.1.2 and P.1-like [17,18]. The mutation profiles of P.1, P.1.1, P.1.2 and P.1-likeII genomes were subjected to covSPECTRUM to evaluate new mutation combinations. The heatmap with dendrogram was constructed for mutation profile visualization using Heatmapper [19].

### Temporal lineage dynamic and phylogenomic analysis

To evaluate the dynamic of SARS-CoV-2 lineages in RS across the epidemic peaks, the relative density profile was accessed using sequenced SARS-CoV-2 genomes combined with two different SARS-CoV-2 datasets available in the EpiCoV database in GISAID as of 18 May 2021: 893 sequences from RS, and 9,113 sequences from other Brazilian regions. In addition, two phylogenomic analysis were performed, one concatenating our dataset of 90 genome sequences with the Nextstrain’ South America dataset (2,961 sequences) and other concatenating only sequences classified as P.1 and P.1-like from Brazil available in GISAID (between December 2019 and April 2021). Analyses were carried out using Nextstrain ncov workflow (https://github.com/) (default parameters), which includes alignment, phylogenetic reconstruction, geographic and ancestral trait reconstruction, and inference of transmission events. Nextstrain uses by default the software IQ-Tree and the substitution model GTR. For the P.1 phylogenomic reconstruction, we adopted a subsampling scheme focused on the RS (group by division, year and month, maintaining 20 sequences per group).

### Data availability

The SARS-CoV-2 sequences were submitted to GISAID and are available for download (**Supplementary Data 1**).

## Results

### Clinical parameters

Of the 90 samples analyzed in this study, 47 were from females (median age 39 years [1– 105 years]), and 43 from males (median age 52 years [6 months–91 years]). Based on available data, 42.2% of the patients (35/83) were hospitalized, and 23.8% (15/63) died. Cough, fever and dyspnea were the main symptoms, whereas cardiopathy and diabetes were the main reported comorbidities (**Supplementary Table 2)**.

### Sequencing data and SARS-CoV-2 lineages diversity

The number of paired-end reads per sequenced genome varied from 94,300 to 697,700, with mean depth coverage ranging from 43x to 558x (**Supplementary Data 1**), and presenting at least 99% coverage breadth to the Wuhan-Hu-1 reference genome (NC_045512.2).

According to Pangolin and to manual verification based on previously described mutational profile [5,6,17,18], our SARS-CoV-2 genomes were assigned to 11 lineages: P.1 (n = 20; 22.2%), B.1.1.28 (n = 19; 21.1%), P.2 (n = 17; 18.9%), B.1.1.33 (n = 15; 16.7%), P.1.2 (n = 7; 7.8%), B.1 (n = 5; 5.6%), B.1.1 (n = 3; 3.3%), B.1.91 (n = 1; 1.1%), A (n = 1; 1.1%), P.1.1 (n = 1; 1.1%) and P.1-likeII (n = 1; 1.1%) (**Supplementary Data 1**). Density profile analysis shows that the main lineages identified in this study are in accordance with the lineage distribution observed for RS and Brazil (**Figure 1**).

Among the first cases of 15 RHCs, we identified lineages A (n = 1; 6.7%), B.1 (n = 5; 33.3%), B.1.1 (n = 2; 13.3%), B.1.1.28 (n = 4; 26.7%), B.1.1.33 (n = 3; 20.0%) (**Figure 2A and 2B**). Notably, lineages A and B.1 were detected in samples from March 2020, whereas the others were identified between March and May 2020 (**Supplementary Data 1**).

**Figure 2.**
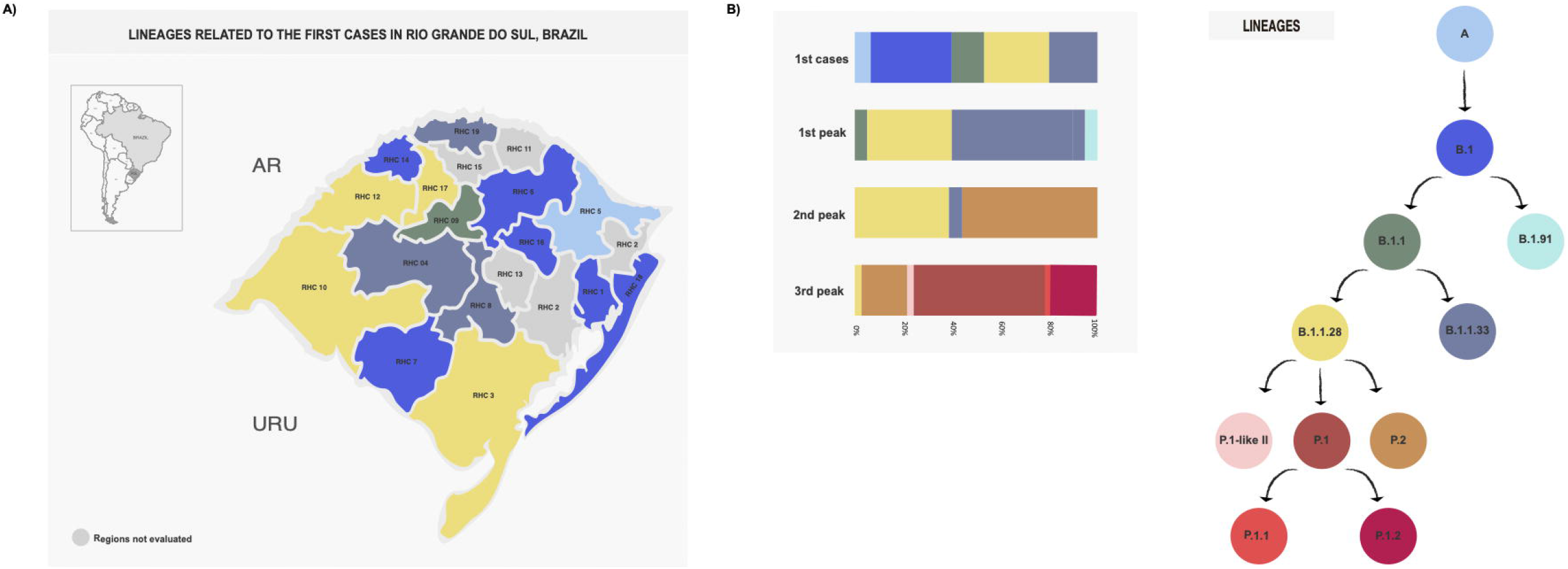
SARS-CoV-2 lineages diversity in Rio Grande do Sul (RS). **A**. Geographical representation of the 19 Regional Health Coordinations, among which 15 are colored according to the SARS-CoV-2 lineage identified in this study. **B**. Profile of SARS-CoV-2 lineages identified in the first cases and in the three epidemic peaks in RS. Moreover, a schematic representation based on lineages’ phylogenetic relationship was used as legend.

Following SARS-CoV-2 introduction in the state, the first peak was marked by lineages B.1 (n = 1; 5.0%), B.1.1.28 (n = 7; 35.0%), B.1.1.33 (n = 11; 55.0%), and B.1.91 (n = 1; 5.0%). In the second peak, P.2 was the most frequent lineage (n = 10; 55.6%), followed by B.1.1.28 (n = 7; 38.9%) and B.1.1.33 (n = 1; 5.6%). In the third peak, P.1 was the most frequent lineage (n = 20; 54.1%), followed by P.1.2 (n = 7; 18.9%), P.2 (n = 7; 18.9%), P.1.1 (n = 1; 2.7%), B.1.1.28 (n = 1; 2.7%) and the recently described P.1-likeII (n = 1; 2.7%) (**Figure 1; Figure 2B**).

### Phylogenomic analysis and mutations

Phylogenetic reconstruction enriched to the South American dataset revealed that the RS genomes grouped according to the Pangolin classification (**Figure 3**). Accordingly, phylogenetic analysis showed that the lineage A sequence from this study grouped with other lineage A sequences from Peru in a major clade containing genomes of lineage A.5 from Bolivia, Uruguay, Chile and Peru. In B.1.1.28 clade, we identified three genomes (85301, 99718 and 22405) grouped with VUI-NP13L, confirmed by the presence of previously reported amino acid change combination [8,12]. Regarding lineage P.2, seven genomes clustered with P.2 sequences from Uruguay, Australia and England.

**Figure 3.**
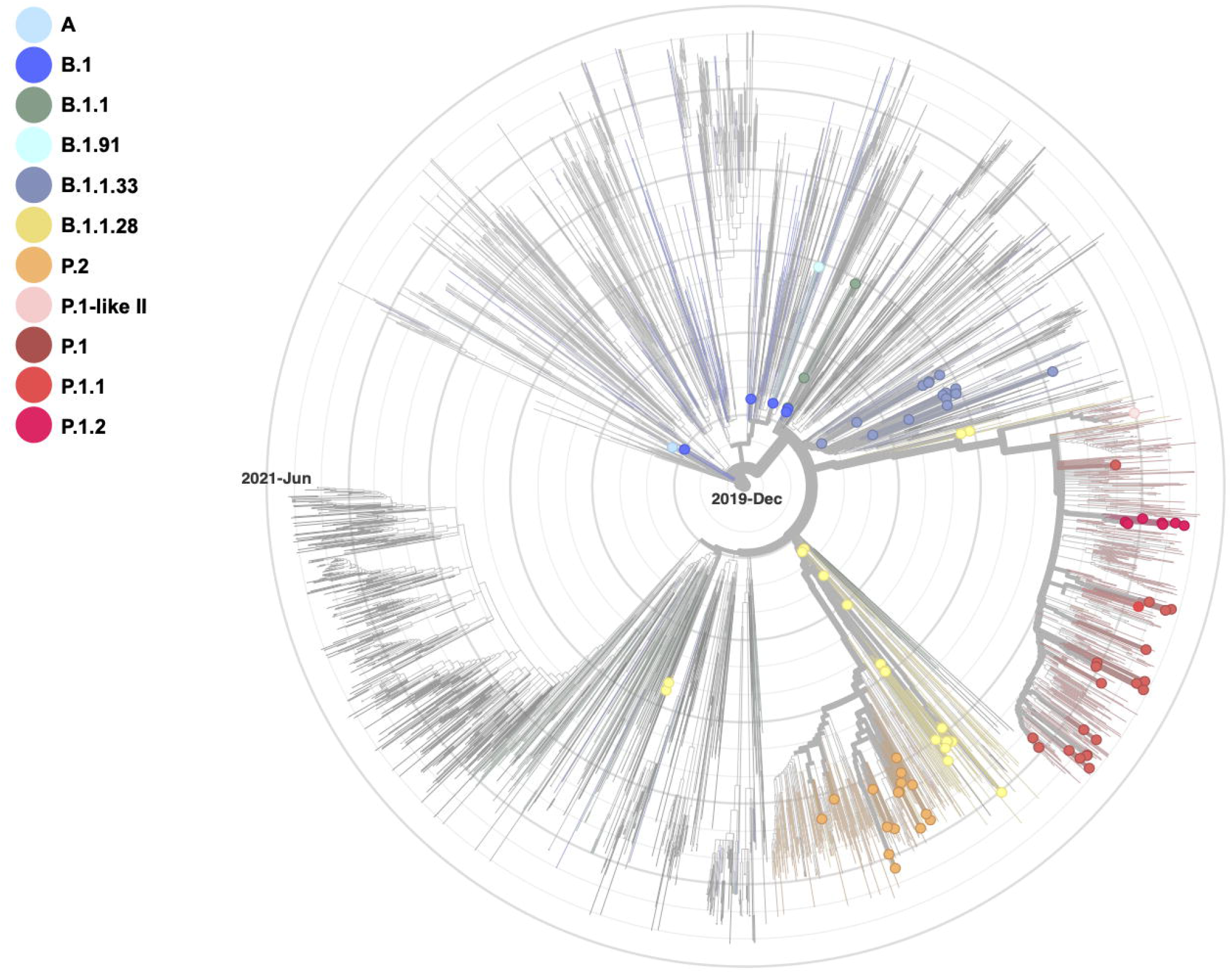
Phylogenomic tree of SARS-CoV-2 genomes. Sequences obtained in this study were combined with a dataset of 2,961 South American genomes. Our SARS-CoV-2 sequences are colored according to lineage classification.

Based on phylogenic tree (**Figure 4**), 29 sequences classified as P.1 and P.1-related were divided in two main groups: one containing sequences of P.1, P.1.1 and P.1.2 (n=28), and other group with the P.1-likeII sequence (n=1). The P.1.1 sequence, recovered in this study, clustered with two sequences from São Paulo, while the P.1.2 sequences grouped in a subcluster containing two other genomes recovered from São Paulo and RS (**Figure 4**).

**Figure 4.**
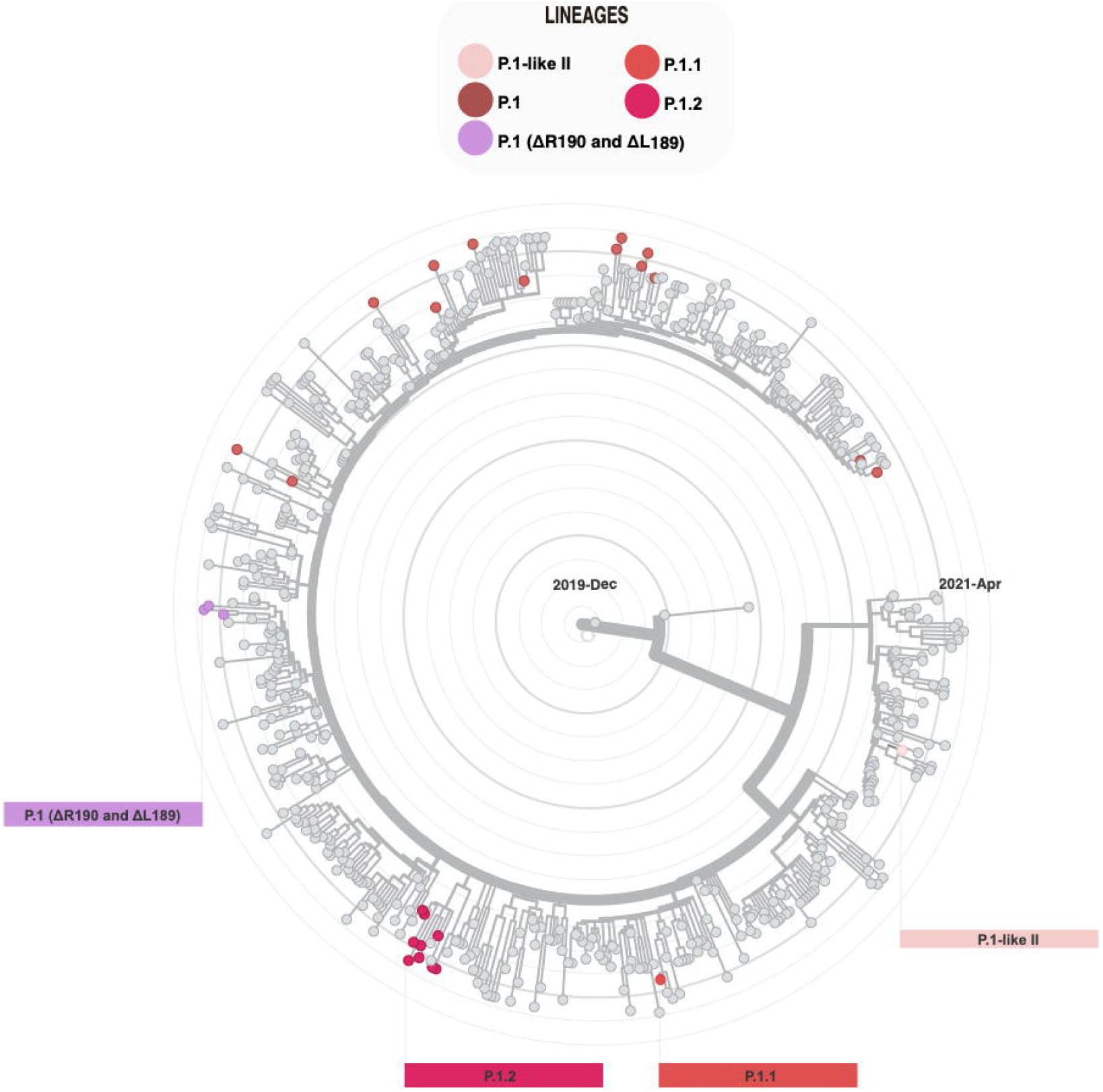
Phylogenomic reconstruction of P.1 and P.1-related lineages. Our SARS-CoV-2 genomes were combined with a dataset of Brazilian P.1, P.1.1, P.1.2 and P.1-like genomes. Sequences recovered in this study are colored according to lineage classification.

Comparing the genomes sequenced in this study with the Wuhan-Hu-1 reference genome (NC_045512.2), a total of 416 nucleotide substitutions, 8 deletions and 1 insertion were found. These mutations comprised 247 amino acid (aa) substitutions and 7 aa deletions: ORF1a (89), ORF1b (47), S (36), ORF3a (21), M (4), ORF6 (6), ORF7a (6), ORF7b (2), ORF8 (13), ORF9b (5), and N (25) (**Supplementary Data 1**).

Regarding P.1 and P.1-related genomes (n = 29), 87 non-synonymous mutations were detected (82 substitutions; 5 deletions), 20 of which are present in all sequences (**Supplementary Table 3**). These mutations were found in ORF1a (32), ORF1b (10), S (22), ORF3a (6), M (1), ORF7a (4), ORF7b (1), ORF8 (3), ORF9b (3) and N (5) (**Figure 5A**). In the Spike gene, the ten lineage-defining mutations (L18F, T20N, P26S, D138Y, R190S, K417T, E484K, N501Y, H655Y and T1027I) were confirmed, and 12 additional mutations/deletions were found: L5F, V1176F, N188S, L189-, R190-, T573I, D614G, E661D, P812S, A845S, T859I, and V1264L. Based on covSPECTRUM analysis, 17 new mutation combination profiles were detected in 18 P.1-related genomes (**Figure 5B**).

**Figure 5.**
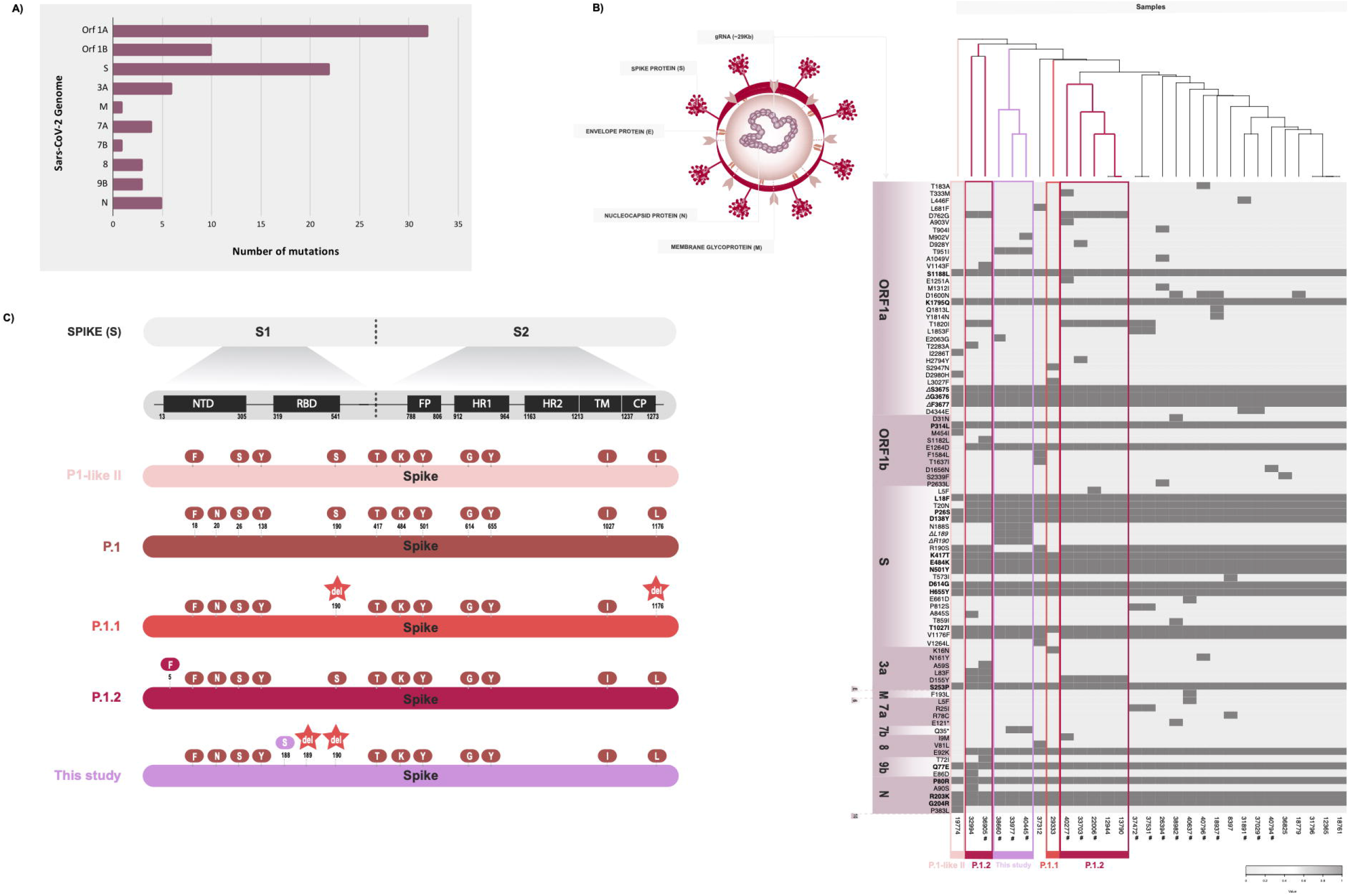
SARS-CoV-2 mutation patterns over time present in P1 and P1-related genomes of this study compared to the reference Wuhan-Hu-1 sequence. **A**. Total of mutations across encoding regions of 29 SARS-CoV-2 genomes of P.1, P.1.1, P.1.2, and P.1-likeII. **B**. Heatmap displaying the non-synonymous mutations present in P.1 and P.1-related sequences. These mutations are plotted according to the respective genome position and are summarized as presence (dark grey) or absence (light grey). A dendrogram based on mutation profile and a schematic representation of SARS-CoV-2 virion structure are also exhibited. **C**. Scheme of mutations on spike proteins highlighting the differences in the protein profile among P.1 and P.1-related lineages.

Three P.1 genomes, recovered from Santo Angelo city in the third peak, showed the mutations ΔL189, ΔR190 and N188S in the Spike, grouping in the heatmap dendrogram (**Figure 5C**). In the phylogenetic analysis, these sequences clustered with other SARS-CoV-2 genomes from Brazil (Bahia, Roraima, Goias and RS) and from Colombia and French Guiana (**Figure 3**; **Figure 4**).

## Discussion

The COVID-19 pandemic has deeply impacted the health system and the economy of Brazil, overcrowding hospitals and overburdening health professionals. Genomic epidemiology has been paramount in understanding SARS-CoV-2 dispersion and to track the evolutive dynamics of viral transmission. In view of this, we investigated the viral dynamics of SARS-CoV-2 in RS from the first cases and across the three main peaks of infection that occurred between July 2020 and April 2021 in the region.

The first individual with SARS-CoV-2 infection confirmed in RS had returned from Italy in early March 2020; our analysis identified it as of lineage B.1, which was also found among other first cases in the state. Traveling records were obtained for three of these introductory cases: two were patients who had been in Europe, whereas the third had been in Rio de Janeiro. Notably, B.1 was the lineage related to the early epidemic outbreak in northern Italy [20,21], hence our findings confirm this route of international introduction of SARS-CoV-2 in RS. B.1.1 lineage, which evolved from B.1 and also circulated in Europe in the beginning of the pandemic, was also found among the first cases in RS.

Interestingly, lineage A was found among the first cases, in a resident of Serafina Correa city who had been in Paraguay; in the phylogenetic analysis, this sample clustered with sequences from Peru. According to Pangolin data, lineage A is considered to be one of the two major lineages of the root of the pandemic, originating in China and spreading to Asia, Europe, Oceania and North America. In South America, lineage A was detected early in the pandemic at low frequencies, with only a few genome sequences reported in Brazil, all from March 2020.

Besides these variants, Brazilian lineages B.1.1.28 and B.1.1.33, which emerged in São Paulo in late February 2020 [22,23], were also detected among the first cases in RS. The dispersion pattern and demographical dynamics of these lineages showed that they were introduced by multiple events, and their spread occurred throughout Brazil by community transmission in early March, and also to other countries [22–25]

Our genomic analyses of the first SARS-CoV-2-confirmed cases in RS suggests that viral introduction in the state was related to both international origin and community-driven transmission. In March 2020, international travel restrictions were adopted to control viral spread, influencing SARS-CoV-2 transmission pattern. Accordingly, previous studies show decreases in imported cases after travel restrictions in Brazil, which, in turn, contribute to increased circulation of local lineages [22,23].

Additionally, our study reveals the temporal frequency and divergence of SARS-CoV-2 lineages in RS over time. As observed in previous Brazilian studies, B.1.1.28 and B.1.1.33 predominated during the first epidemic peak (July and August 2020) [7,8,12,22]. Selection of these lineages over others might be explained by mutations associated with higher viral fitness and severe disease [26]. Moreover, the V1176F mutation in B.1.1.28 is predicted to increase the flexibility of the stalk domain in the Spike protein trimmer, facilitating its binding to the ACE2 receptor [27].

B.1.1.28 and B1.1.33 continued to circulate in the second peak; however, an emergence of P.2 was observed. This lineage descends from B.1.1.28, carrying the lineage-defining mutations ORF1ab:L3468V, ORF1ab:synC11824U, N:A119S, and S:E484K [7]. According to our previous study, P.2 has been massively circulating in RS since October 2020 [12]. Additionally, in the present study three genomes (two in the second peak and one in third peak) classified as B.1.1.28 were shown to be VUI-NP13L, a potential new lineage [8,12]. VUI-NP13L was first reported in RS in August 2020, disseminating afterwards [12], and our results reinforce that this lineage continued to circulate at low frequency in RS.

The third peak was marked by P.1 and P.2 as the predominant lineages. Both descend from B.1.1.28 have distinct origins, and share the S:E484K mutation [28]. P.1 was first identified in four travelers arriving in Japan from Amazonas, Brazil (on 2 January 2021) [29], and was later confirmed in several Amazonas samples (in November 2020) [5]. In our study, P.1 and P.1-related lineages were the most frequent in RS since the first week of February 2021, becoming the dominant lineages in the third peak; P.1 was first identified in late January 2021 in Gramado, a touristic town that annually receives 6.5 million visitors [30]. The P.1 rapid dispersion might be related to its higher transmission rate [31], higher viral loads [6], greater affinity to ACE2 receptor [32], and the ability to resist neutralizing antibodies, either from natural infections or vaccine-induced [28,32,33]. Moreover, P.1 spread may also be influenced by relaxed social distancing measures at holidays and vacation, especially during the summer period in the state.

P.1 lineage became a VOC based on lineage-defining mutations in proteins S (L18F, T20N, P26S, D138Y, R190S, K417T, E484K, N501Y, H655Y and T1027I), NSP3 (S370L and K977Q), NSP13 (E341D), NS8 (E92K) and N (P80R) [6,17]. The high number of lineage-defining mutations for P.1 is considered a result of sequential infection steps during B.1.1.28 evolution in Amazonas [17]. Our results show the expressive spread of P.1 across Brazilian states and the high genetic diversity of P.1-related lineages in RS since February 2021. Besides lineage-defining mutations, 87 additional mutations were found among our P.1 genomes. Moreover, 17 new mutation combinations were observed in 18 genomes, suggesting that independent events are related to the observed variants. Accordingly, the P.1 genomes obtained here are present in several clades in the phylogenomic tree; additionally, this rapid diversification was evidenced by P.1.1 and P.1.2 detection.

Among our P.1 sequences, three genomes displayed two deletions in the S gene (ΔL189 and ΔR190) in the N-terminal domain (NTD), which were recently found in four other genomes recovered in Brazil [34]. Deletions in NTD have been reported during prolonged infection of immunocompromised patients [35,36] and subsequent transmission [37]. McCarthy et al. [35] observed deletions in the S gene, of which >97% maintained the open reading frame, and 90% occurred in four sites, named “recurrent deletion regions 1–4” (RDRs). Considering these RDRs, the ΔL189 and ΔR190 deletions are located between RDR2 (position 139–146) and RDR3 (position 210–212). Although both deletions were not reported to affect the NTD antigenic-supersite, they might lead to conformational changes in exterior loops, affecting antibody binding outside the antigenic-supersite [34]. Deletions in NTD have been associated with resistance to antibody neutralization, suggesting an improvement in virus fitness by evading the host’s immune response, an evolutive process due to immune pressure [34–36].

Our results allowed the analysis of the first SARS-CoV-2 lineages and of the main epidemic peaks in different geographic regions in RS, despite the limited number of investigated samples. Furthermore, the analysis of SARS-CoV-2 diversity from three epidemic peaks provided a profile of circulating lineages in the State and in Brazil and highlighted the massive spread of P.1 and P.1-related lineages in RS. Hence, this study contributes to the genomic surveillance of SARS-CoV-2 and reinforces its importance to guide health authorities in decisions for the control of COVID-19.

## Author contributions

APMV, JP and FQM: design of the study, generating sequences, data analysis, writing the manuscript; RS and FHS: analyzing sequences and data analysis; TSG: concept of the study, collection of samples, reviewing the manuscript; LGM: data acquisition; AS: design of the study and reviewing the manuscript; ABGV: concept of the study, data acquisition and analysis, reviewing draft and writing the manuscript. All authors critically revised the manuscript, approved the final version to be published, and agreed to be accountable for all aspects of the work.

## Acknowledgments

We thank Dr. Vanessa Mattevi, Dr. Marília Zandoná, and Ludmila Fiorenzano Baethgen for providing technical support. We thank all the authors who have shared genome data on GISAID, and we have included a table (**Supplementary Data 1**) listing the authors and institutes involved.

## Supplementary Material

**Supplementary Data 1:** Metadata of the 90 SARS-CoV-2 genomes recovered in this study.

**Supplementary Table 1**. Data of the first confirmed COVID-19 cases in RHCs evaluated in the present study.

**Supplementary Table 2**. Characteristics of the SARS-CoV-2 cases analyzed in this study.

**Supplementary Table 3**. Mutations detected in P.1 (n = 20), P.1.1 (n = 1), P.1.2 (n = 7) and P.1-likeII (n = 1) in the present study. A total of 87 nonsynonymous substitutions are shown.

