## Supplemental Table 1 for "SARS-CoV-2 introduction and lineage dynamics across three epidemic peaks in Southern Brazil: massive spread of P.1"

**Supplementary Table 1**. Data of the first confirmed COVID-19 cases in RHCs evaluated in the present study.

| **First symptom** | **RHC** | **Lineage** | **Sex** | | **Age^a^** | **Symptoms** | **Comorbidity** | | | **Hosp** | | **Mech Vent** | | **Outcome** | | **Traveling (2020)** |
| --- | --- | --- | --- | --- | --- | --- | --- | --- | --- | --- | --- | --- | --- | --- | --- | --- |
| 29/02/20 | 1 | B.1 | Male | | 60 | Cg, Fv, ST | None | | | No | | No | | Cure | | Italy (Feb16-Feb23) |
| 08/03/20 | 14 | B.1 | Male | | 31 | An, Fv, ST | N.I. | | | No | | No | | Cure | | Lives in Ireland, was in Brazil visiting parents |
| 11/03/20 | 7 | B.1 | Male | | 59 | Cg, Dy, Fv, O_2_, RD, ST | Cardiopathy | | | Yes | | Yes | | Cure | | Rio de Janeiro (Mar1) |
| 14/03/20 | 5 | A | Male | | 63 | Cg, Dy, Fv, RD, ST, Vo | DM | | | Yes | | Yes | | Deah | | Paraguay (Mar12) |
| 15/03/20 | 16 | B.1 | Male | | 57 | Cg, Cz, Fv, ST | N.I. | | | No | | No | | Cure | | Madrid, Zurich, Saint Mortiz (Mar3-Mar15) |
| 16/03/20 | 4 | B.1.1.33 | Male | | 29 | An, Cg, Di, Fv, RD, ST | N.I. | | | No | | No | | Cure | | None |
| 16/03/20 | 10 | B.1.1.28 | Male | | 55 | Cg, Fv | N.I. | | | No | | No | | Cure | | None |
| 17/03/20 | 3 | B.1.1.28 | Male | | 54 | Cg, Fv, O_2_, ST | Neoplasia | | | Yes | | No | | Cure | | None |
| 17/03/20 | 6 | B.1.1 | Male | | 29 | Fv, Di, Dy, He, Na, Vo | N.I. | | | No | | No | |  | | None |
| 18/03/20 | 18 | B.1 | Male | | 28 | Ch, Cg, Fv, ST | N.I. | | | No | | No | | Cure | | Trip to France, with symptoms on the way back |
| 22/04/20 | 8 | B.1.1.33 | Male | | 65 | Cg, Dy, Fv, O_2_, RD | Cardiopathy, DM | | | Yes | | Yes | | Cure | | Contact with infected person from Lajeado (RHC 16) |
| 23/04/20 | 12 | B.1.1.28 | Male | | 39 | Cg, Fe, O_2_, RD, Vo | Obesity | | | Yes | | Yes | | Death | | None |
| 25/04/20 | 9 | B.1.1 | Female | | 29 | BP, Cg, Cz, Fv | N.I. | | | No | | No | | Cure | | Carazinho (Apr22, RHC 6) |
| 13/05/20 | 19 | B.1.1.33 | Female | | 31 | Cg, Dy, Fv, O_2_, RD | None | | | Yes | | Yes | | Death | | None |
| 31/05/20 | 17 | B.1.1.28 | Female | | 53 | Ar, Cg, Fv, He, My, RD, ST | N.I. | | | No | | No | |  | | None |
| An, anosmia; Ar, arthralgia; BP, back pain; Ch, chills; Cg, cough; Cz, coryza; Di, diarrhea; DM, Diabetes mellitus; Dy, dyspnea; Fv, fever; He, headache; Hospit, hospitalization; Mech vent, mechanical ventilation; My, mylagia; Na, nausea; N.I., not informed; O_2_, oxygen saturation <95%; RD, respiratory distress; RHC, Regional Health Coordination; ST, sore throat; Vo, vomiting. | | | | | | | | | | | | | | | | |
| ^a^Age in years. | |  | |  |  |  | |  |  | |  | |  | |  | |
