## Supplemental Table 2 for "SARS-CoV-2 introduction and lineage dynamics across three epidemic peaks in Southern Brazil: massive spread of P.1"

**Supplementary Table 2.** Characteristics of the SARS-CoV-2 cases analyzed in the study.

| **Variable** | | **N** | **%** |
| --- | --- | --- | --- |
| Sex | |  |  |
|  | Male | 43 | 47.78 |
|  | Female | 47 | 52.22 |
| Age (years) | |  |  |
|  | 0–9 | 6 | 6.67 |
|  | 10–19 | 4 | 4.44 |
|  | 20–39 | 30 | 33.33 |
|  | 40–59 | 25 | 27.78 |
|  | 60–79 | 18 | 20.00 |
|  | ≥80 | 7 | 7.78 |
| Symptom^a^ | |  |  |
|  | Cough | 57/82 | 69.51 |
|  | Fever | 51/84 | 60.71 |
|  | Dyspnea | 35/70 | 50.00 |
|  | Respiratory distress | 30/71 | 42.25 |
|  | O_2_ <95% | 28/69 | 40.58 |
|  | Headache | 25/71 | 35.21 |
|  | Sore throat | 28/80 | 35.00 |
|  | Fatigue | 16/66 | 24.24 |
|  | Anosmia | 16/66 | 24.24 |
|  | Diarrhea | 14/70 | 20.00 |
|  | Myalgia | 14/71 | 19.72 |
|  | Vomiting | 10/67 | 14.92 |
|  | Coryza | 10/71 | 14.08 |
|  | Ageusia | 9/64 | 14.06 |
|  | Abdominal pain | 5/63 | 7.94 |
|  | Other^b^ | 10/71 | 14.08 |
| Underlying condition^a^ | |  |  |
|  | Cardiopathy | 17/74 | 22.97 |
|  | Diabetes | 16/74 | 21.62 |
|  | Obesity | 6/74 | 8.11 |
|  | Hypertension | 5/74 | 6.76 |
|  | Pregnant | 4/74 | 5.40 |
|  | Neurological disease | 3/74 | 4.05 |
|  | Cancer | 3/74 | 4.05 |
|  | Pneumopathy | 3/74 | 4.05 |
|  | Other^c^ | 9/74 | 12.16 |
| Hospitalization^a^ | | 35/83 | 42.17 |
| Mechanical ventilation | | 14/81 | 17.28 |
| Outcome^a^ | |  |  |
|  | Cure | 48/63 | 76.19 |
|  | Fatality | 15/63 | 23.81 |
| Lineage^d^ | |  |  |
|  | A | 1 | 1.11 |
|  | B.1 | 5 | 5.55 |
|  | B.1.1 | 3 | 3.33 |
|  | B.1.91 | 1 | 1.11 |
|  | B.1.1.28 | 19 | 21.11 |
|  | B.1.1.33 | 15 | 16.67 |
|  | P.2 | 17 | 18.89 |
|  | P.1-likeII | 1 | 1.11 |
|  | P.1 | 21 | 23.33 |
|  | P.1.2 | 7 | 7.78 |
| N, number of cases | |  |  |
| ^a^Some cases with no information; shown are the number of cases presenting the symptom in relation to the number of cases with available information for that symptom. | | | |
| ^b^Other symptoms include nasal obstruction (2/71), back pain (2/71), arthralgia (1/71), chills (1/71), loss of appetite (1/71), nausea (1/71). | | | |
| ^c^Other underlying conditions include asthma (2/74), post-partum (2/74), nephropathy (1/74), smoking (1/74), alcoholist (1/74), immunodeficiency (1/74), immunologic disease (1/74). | | | |
| ^d^SARS-CoV-2 lineages shown in the order they were detected in RS. | | | |
