## Supplemental Table 3 for "SARS-CoV-2 introduction and lineage dynamics across three epidemic peaks in Southern Brazil: massive spread of P.1"

**Supplementary Table 3.** Mutations detected in P.1 (n = 20), P.1.1 (n = 1), P.1.2 (n = 7) and P.1-likeII (n = 1) in the present study. A total of 87 nonsynonymous substitutions are shown.

| Gene | Effect | Mutation | Lineages | Number of occurrences | Frequency (%) |
| --- | --- | --- | --- | --- | --- |
| ORF1a | Substitution | T183A | P.1 | 1 | 3.4 |
|  |  | T333M | P.1.2 | 1 | 3.4 |
|  |  | L446F | P.1.1 | 1 | 3.4 |
|  |  | L681F | P.1 | 1 | 3.4 |
|  |  | D762G | P.1.2 | 7 | 24.1 |
|  |  | A903V | P.1.2 | 1 | 3.4 |
|  |  | T904I | P.1 | 1 | 3.4 |
|  |  | M902V | P.1 | 1 | 3.4 |
|  |  | D928Y | P.1.2 | 1 | 3.4 |
|  |  | T951I | P.1 | 3 | 10.3 |
|  |  | A1049V | P.1 | 1 | 3.4 |
|  |  | V1143F | P.1.2 | 1 | 3.4 |
|  |  | S1188L | P.1, P.1-likeII, P.1.1, P.1.2 | 29 | 100.0 |
|  |  | E1251A | P.1.2 | 1 | 3.4 |
|  |  | M1312I | P.1 | 1 | 3.4 |
|  |  | D1600N | P.1 | 4 | 13.8 |
|  |  | K1795Q | P.1, P.1-likeII, P.1.1, P.1.2 | 29 | 100.0 |
|  |  | Q1813L | P.1 | 1 | 3.4 |
|  |  | Y1814N | P.1 | 1 | 3.4 |
|  |  | T1820I | P.1, P.1.2 | 9 | 31.0 |
|  |  | L1853F | P.1 | 2 | 6.9 |
|  |  | E2063G | P.1 | 1 | 3.4 |
|  |  | T2283A | P.1.2 | 1 | 3.4 |
|  |  | I2286T | P.1-likeII | 1 | 3.4 |
|  |  | H2794Y | P.1.2 | 1 | 3.4 |
|  |  | S2947N | P.1.1 | 1 | 3.4 |
|  |  | D2980H | P.1-likeII | 1 | 3.4 |
|  |  | L3027F | P.1.1 | 1 | 3.4 |
|  |  | D4344E | P.1 | 2 | 6.9 |
|  | Deletion | S3675- | P.1, P.1-likeII, P.1.1, P.1.2 | 29 | 100.0 |
|  |  | G3676- | P.1, P.1-likeII, P.1.1, P.1.2 | 29 | 100.0 |
|  |  | F3677- | P.1, P.1-likeII, P.1.1, P.1.2 | 29 | 100.0 |
| ORF1b | Substitution | D31N | P.1 | 1 | 3.4 |
|  |  | P314L | P.1, P.1-likeII, P.1.1, P.1.2 | 29 | 100.0 |
|  |  | M454I | P.1.2 | 1 | 3.4 |
|  |  | S1182L | P.1.2 | 1 | 3.4 |
|  |  | E1264D | P.1, P.1.1, P.1.2 | 28 | 96.6 |
|  |  | F1584L | P.1 | 1 | 3.4 |
|  |  | T1637I | P.1 | 1 | 3.4 |
|  |  | D1656N | P.1 | 1 | 3.4 |
|  |  | S2339F | P.1 | 1 | 3.4 |
|  |  | P2633L | P.1 | 1 | 3.4 |
| S | Substitution | L5F | P.1.2 | 1 | 3.4 |
|  |  | L18F^a^ | P.1, P.1-likeII, P.1.1, P.1.2 | 29 | 100.0 |
|  |  | T20N^a^ | P.1, P.1.1, P.1.2 | 28 | 96.6 |
|  |  | P26S^a^ | P.1, P.1-likeII, P.1.1, P.1.2 | 29 | 100.0 |
|  |  | D138Y^a^ | P.1, P.1-likeII, P.1.1, P.1.2 | 29 | 100.0 |
|  |  | N188S | P.1 | 3 | 10.3 |
|  |  | R190S^a^ | P.1, P.1-likeII, P.1.2 | 25 | 86.2 |
|  |  | K417T^a^ | P.1, P.1-likeII, P.1.1, P.1.2 | 29 | 100.0 |
|  |  | E484K^a^ | P.1, P.1-likeII, P.1.1, P.1.2 | 29 | 100.0 |
|  |  | N501Y^a^ | P.1, P.1-likeII, P.1.1, P.1.2 | 29 | 100.0 |
|  |  | T573I | P.1 | 1 | 3.4 |
|  |  | D614G | P.1, P.1-likeII, P.1.1, P.1.2 | 29 | 100.0 |
|  |  | H655Y^a^ | P.1, P.1-likeII, P.1.1, P.1.2 | 29 | 100.0 |
|  |  | E661D | P.1 | 1 | 3.4 |
|  |  | P812S | P.1 | 2 | 6.9 |
|  |  | A845S | P.1.2 | 1 | 3.4 |
|  |  | T859I | P.1 | 1 | 3.4 |
|  |  | T1027I^a^ | P.1, P.1-likeII, P.1.1, P.1.2 | 29 | 100.0 |
|  |  | V1176F | P.1, P.1-likeII, P.1.2 | 28 | 96.6 |
|  |  | V1264L | P.1 | 1 | 3.4 |
|  | Deletion | L189- | P.1 | 3 | 10.3 |
|  |  | R190- | P.1 | 3 | 10.3 |
| ORF3a | Substitution | K16N | P.1.1 | 1 | 3.4 |
|  |  | N161Y | P.1 | 1 | 3.4 |
|  |  | A59S | P.1.2 | 1 | 3.4 |
|  |  | L83F | P.1.2 | 2 | 6.9 |
|  |  | D155Y | P.1.2 | 7 | 24.1 |
|  |  | S253P | P.1, P.1-likeII, P.1.1, P.1.2 | 29 | 100.0 |
| M | Substitution | F193L | P.1 | 1 | 3.4 |
| ORF7a | Substitution | L5F | P.1 | 1 | 3.4 |
|  |  | R25I | P.1 | 2 | 6.9 |
|  |  | R78C | P.1 | 1 | 3.4 |
|  | Stop codon | E121* | P.1 | 1 | 3.4 |
| ORF7b | Stop codon | Q35* | P.1 | 2 | 6.9 |
| ORF8 | Substitution | I9M | P.1.2 | 1 | 3.4 |
|  |  | V81L | P.1 | 1 | 3.4 |
|  |  | E92K | P.1, P.1.1, P.1.2 | 28 | 96.6 |
| ORF9b | Substitution | T72I | P.1.2 | 1 | 3.4 |
|  |  | Q77E | P.1, P.1-likeII, P.1.1, P.1.2 | 29 | 100.0 |
|  |  | E86D | P.1.2 | 1 | 3.4 |
| N | Substitution | P80R | P.1, P.1-likeII, P.1.1, P.1.2 | 29 | 100.0 |
|  |  | A90S | P.1.2 | 1 | 3.4 |
|  |  | R203K | P.1, P.1-likeII, P.1.1, P.1.2 | 29 | 100.0 |
|  |  | G204R | P.1, P.1-likeII, P.1.1, P.1.2 | 29 | 100.0 |
|  |  | P383L | P.1-likeII | 1 | 3.4 |

^a^P.1 lineage defining mutations.

* No amino acid (stop codon).
